# Targeting the energy metabolism of melanoma cells: FX-11 acts as a mitochondrial uncoupler

**DOI:** 10.1101/2024.11.04.621801

**Authors:** Jasmin Renate Preis, Catherine Rolvering, Mélanie Kirchmeyer, Iris Behrmann, Claude Haan

**Affiliations:** Department of Life Sciences and Medicine, University of Luxembourg, Belvaux, Luxembourg

**Keywords:** melanoma, metabolism, FX-11, mitochondrial uncoupling, AMPK, mTOR, lactate dehydrogenase inhibitor

## Abstract

In a compound screen on melanoma cells, which included different microenvironmental contexts aiming at better representing the environmental and growth characteristics of tumours, we identified several drugs efficiently suppressing cell viability. Here we focus on results obtained with FX-11, reported to be an inhibitor of lactate dehydrogenase (LDH). FX-11 dose-dependently inhibited growth of 624Mel and Wm3248 melanoma cells grown in a modular physiologic medium (MPM). Importantly, FX-11 was able to reduce the growth of the corresponding drug-resistant melanoma sublines equipotently.

When testing the on-target activity of FX-11, the results were unexpected: in contrast to other LDH inhibitors (used as positive controls), FX-11 did not decrease the NAD^+^/NADH ratio, the glucose uptake, or the lactate secretion of melanoma cells. However, in seahorse assays, FX-11 increased the oxygen consumption rate as well as the extracellular acidification rate of cells, behaving like the mitochondrial uncouplers FCCP and Bam15. Treatment with FX-11, FCCP, or Bam15 induced an increase in acetyl-CoA carboxylase phosphorylation and a decrease in serine 65 phosphorylation of the eukaryotic initiation factor 4E-binding protein 1, indicative of AMPK activation by a decreased ATP/ADP ratio. Importantly, FX-11 and Bam15 drastically decreased the mitochondrial membrane potential in contrast to the LDH inhibitors LDH-IN-I and GNE-140.

Taken together, we provide evidence that FX-11 effectively inhibits the growth of melanoma cells, including drug-resistant ones. FX-11 profoundly affects their energy metabolism, although it does not seem to act as other LDH inhibitors, but as a mitochondrial uncoupler.

## Introduction

Melanoma is an aggressive form of skin cancer, and its incidence has been rising over the last years (Michielin et al., 2019). Metastatic melanoma is associated with a poor overall survival. Around half of melanoma patients harbour a mutation in *BRAF*, leading to constitutive MAPK signalling (Ascierto et al., 2012). The development of small molecule inhibitors targeting the BRAF kinase and its downstream target MEK significantly extended the survival of melanoma patients. However, resistance to the treatment inevitably develops and patients experience disease relapse (Schadendorf et al., 2018). Different resistance mechanisms have been described; besides MAPK-dependent and -independent mechanisms, also metabolic rewiring has been observed (Kozar et al., 2019). The constitutive activation of the MAPK pathway drives the “Warburg effect” in untreated melanoma. Treatment with MAPK inhibitors initially decreases glucose conversion to lactate and increases mitochondrial metabolism. In treatment-resistant melanoma cells, the reactivation of the MAPK pathway is described to result in restored glycolysis with a concomitant higher mitochondrial metabolism (Alkaraki et al., 2021).

Most studies of metabolic rewiring *in vitro* are performed in standard cell culture media with non-physiological concentration of metabolites. Only in recent years, more physiologically relevant media, better reflecting the plasma concentration of metabolites, have been developed (Ackermann & Tardito, 2019; Cantor, 2019). First studies have highlighted differences in metabolic reactions of cancer cells, depending on the metabolite concentrations present in the culture medium, with important consequences for drug discovery (Cantor et al., 2017; Muir & Vander Heiden, 2018; Rossiter et al., 2021; Vande Voorde et al., 2019). We implemented a physiological medium (MPM – modular physiological medium) (Preis, Rolvering, Kirchmeyer, Halilovic, et al., 2024, Preprint) based on previous published media formulations and on published plasma metabolite concentrations (Cantor et al., 2017; Psychogios et al., 2011; Vande Voorde et al., 2019). We used this medium to screen a metabolism-oriented compound library in different cellular contexts (2D culture under “normoxia” or hypoxia and 3D spheroid cultures) and identified different compounds, amongst them FX-11, which inhibited melanoma cell growth (Preis, Rolvering, Kirchmeyer, Kreis, et al., 2024, Preprint).

In this study we further investigated the effects of FX-11 on drug-naïve and derived Encorafenib/Binimetinib-resistant 624Mel and Wm3248 melanoma lines (Preis, Rolvering, Kirchmeyer, Kreis, et al., 2024, Preprint) in MPM. FX-11 was described to be a selective NADH-competitive small molecule inhibitor of lactate dehydrogenase (LDH), and several studies have reported its efficacy in different types of cancer (Hou et al., 2021; Le et al., 2010; Rellinger et al., 2017). In our hands, FX-11 dose-dependently inhibited growth of drug-sensitive and drug-resistant melanoma cells in MPM under all tested culture conditions. Surprisingly, we find that FX- 11 did not decrease the NAD^+^/NADH ratio, glucose uptake, or lactate secretion of cells, as did other LDH inhibitors we used as controls. However, FX-11 behaved like the mitochondrial uncouplers FCCP and Bam15 in Seahorse assays. Importantly, FX-11 and Bam15 very efficiently decreased the mitochondrial membrane potential, in contrast to the LDH inhibitors.

Taken together, we analysed the effects of FX-11 on several melanoma cell lines in physiologic medium, and we find FX-11 not to recapitulate the features of LDH inhibitors, but to rather act as a mitochondrial uncoupler.

## Materials and methods

### Cell culture and materials

All cells were grown at 37 °C in a water-saturated atmosphere at 5% CO_2_. The melanoma cell lines (624Mel (Dr. Ruth Halaban) and Wm3248 (Rockland)) were maintained in RPMI-1640 Glutamax™ (Gibco) supplemented with 10% FBS and 1% penicillin/streptomycin. If not indicated otherwise, cells were cultured in the in-house physiologic medium MPM for all experiments (Preis, Rolvering, Kirchmeyer, Halilovic, et al., 2024, Preprint), supplemented with 2.5% dialyzed FBS. FX-11 (Calbiochem, Sigma-Aldrich), LDH-IN-1 (MedChemExpress), (R)-GNE140 (MedChemExpress), FCCP (Cayman Chemical), and Bam15 (MedChemExpress) were dissolved in DMSO, aliquoted and stored at −20°C prior to use. Multiple freeze/thaw cycles were avoided due to potential loss of activity of the compounds.

### Generation of drug-resistant (DR) melanoma cell lines

Drug-resistant (DR) melanoma cells were generated from parental cell lines by long-term culture under continuous presence of BRAF inhibitor Encorafenib and MEK inhibitor Binimetinib, corresponding to approximatively ten times the IC_50_ concentrations (Preis, Rolvering, Kirchmeyer, Kreis, et al., 2024, Preprint). Drug-naïve and drug-resistant cell lines were authenticated by STR profiling. Resistant cell lines were maintained under continuous exposure to Encorafenib and Binimetinib at concentrations used to generate the resistant lines (for 624MelDR: 30nM Encorafenib and 400nM Binimetinib; for Wm3248DR: 15nM Encorafenib, 200nM Binimetinib).

### 2D cell proliferation assays

Cells were seeded in 96-well black-walled µClear plates (Greiner) in MPM for 2D adherent cell growth. The inhibitors were added as indicated in the figures. At the experiment endpoint at 72h, cell viability was measured with two different readouts. The CyQuant Direct Cell Proliferation Assay Kit (ThermoFisher Scientific, λ_ex_: 489 ± 12 nm/λ_em_: 530 ± 10 nm) and PrestoBlue Cell Viability Reagent (Thermofisher Scientific; λ_ex_: 547 ± 12 nm/λ_em_: 612 ± 20 nm) were used in a duplex assay and fluorescence was read on a Cytation 5 Cell Imaging Multi-Mode Reader (Agilent Technologies). GraphPad Prism v9 software was used to calculate the half-maximal inhibitor concentration values (IC_50_) using the nonlinear log (inhibitor) *vs*. response-variable slope (four parameter) equation. Dose-response curves for FX-11 were constrained to 0. Only curve fittings with an R^2^ value above 0.95 were considered.

### 2D live/dead cell assay upon drug treatment

Cells were seeded in 96-well flat-bottom plates (Greiner Bio-one) for 2D adherent cell growth. Treatments were applied as described in the figure legends. After the times indicated in the figure legends, cells were stained with Hoechst 33342 (1µg/mL; nuclear live cell stain, Life Technologies) and Sytox Orange (1µM; nuclear dead cell stain, Life Technologies) for 30min and imaged on a Cytation 5 Cell Imaging Multi-Mode Reader (Agilent Technologies) using DAPI and RFP filters/LED cubes. The Gen5 software (Agilent Technologies) was used to determine total cell numbers (Hoechst-stained nuclei) and cell death (Sytox orange-positive nuclei).

### Apoptosis assay

Apoptosis was monitored by measuring caspase-3 activity through cleavage of the AC-DEVD- AFC peptide (AlfaAesar) and release of fluorescent 7-amino-4-trifluoromethylcoumarin (AFC) in solution. Briefly, melanoma cells were seeded in µClear plates (Greiner) with or without treatment. After 16 or 24h, cells were lysed with 3x ReLy buffer (150mM Tris (pH 7.4), 300mM NaCl, 30% glycerol, 1% Triton-X, 0.3% CHAPS, 6mM EDTA (pH 8.0), 6mM DTT, 75μM Ac-DEVD-AFC), of which DTT and Ac-DEVD-AFC were added just prior to use. Fluorescence of cleaved AFC (λ_ex_: 400 ± 15 nm/λ_em_: 505 ± 20 nm) was measured 2h after addition of the reagents, using an Cytation 5 Cell Imaging Multi-Mode Reader (Agilent Technologies). 2µM Staurosporine (Santa Cruz Biotechnology) or 30µM TG101348 (Selleck chemicals) were included as positive controls. Encorafenib, which induces growth reduction without apoptosis induction, was added as a negative control.

### ROS detection using flow cytometry

The generation of reactive oxygen species (ROS) was quantified with the fluorogenic probe CM- H2DCFDA (6-chloromethyl-2’,7’-dichlorodihydrofluorescein diacetate, acetyl ester) (Thermo Fisher Scientific). Cells were treated with inhibitors for 4h, before detachment and staining with 5µM CM-H2DCFDA for 30min. After washing steps, DCF (dichlorofluorescein) fluorescence was assessed in the FITC (fluorescein isothiocyanate) channel of a FACS Canto flow cytometer (BD Bioscience). Dead cell populations were excluded by staining with the Live/Dead Fixable Near-IR cell stain (Thermo Fisher Scientific). Analysis was performed with the FlowJo v10.8.1 software.

### Determination of the NAD^+^/NADH ratio

Cells were seeded at 10,000 cells per well in 96-well plates and permitted to adhere. Inhibitors were added and, following a 24h incubation, NAD^+^ and NADH was quantified using the NAD/NADH-Glo Assay kit (Promega, G9072) according to the manufacturer’s instructions. Luminescence was measured in a Cytation 5 plate reader (Agilent Technologies).

### YSI measurements to determine glucose and lactate concentrations

624Mel cells were seeded in 6-well plates at 150,000 cells per well in 2mL MPM medium. After attachment of cells, the medium was exchanged for fresh or drug-containing medium. After 24h, medium was collected for metabolite quantification and cells were counted for each condition. Collected medium samples were filtered (Phenex-RC 4mm Syringe Filters 0.2µm) to remove particles prior measurement. Quantitative values for glucose and lactate concentrations were acquired using the YSI 2950D Biochemistry Analyzer (YSI incorporated). To calculate consumption (c) and secretion (s) rates, the medium control value was subtracted from the absolute metabolite concentrations and normalized to the cell count according to the following equation, adapted from Vande Voorde et al., 2019:

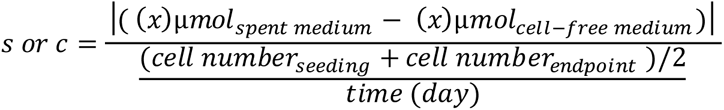

### Analyses using the Seahorse XFe96 analyser

A Seahorse XFe96 analyser (Agilent Technologies) was used to measure the Oxygen Consumption Rate (OCR) and the Extracellular Acidification Rate (ECAR). One day before the assay, melanoma cells were seeded in poly-D-lysine-coated Seahorse 96-well plates, at a density of 10,000 cells/well for 624Mel and 20,000 cells/well for Wm3248 cells, in RPMI 10% FBS and 1% penicillin/streptomycin. On the day of the assay, cells were washed with Seahorse XF RPMI (pH 7.4) supplemented with 10mM glucose, 2mM glutamine, and 1mM pyruvate, and incubated in a non-CO_2_ incubator at 37°C for 1h. To examine the effects of FX-11 on bioenergetics, various concentrations of the test compound were injected and OCR and ECAR was recorded for 1h. For the XF Cell Mito Stress Test, sequential injections of 1.5µM Oligomycin (ATP synthase inhibitor), mitochondrial uncouplers (FCCP, Bam15, or, for comparison, FX-11), and 0.5µM Rotenone and Antimycin A (complex I and complex III inhibitors, respectively) were performed. The OCR and ECAR values were normalized to cell counts determined with 1µg/mL Hoechst 33342 (Life Technologies) staining and imaging on a Cytation 5 Cell Imaging Multi-Mode Reader (Agilent Technologies).

Technical replicates were excluded if drug injection in the Seahorse failed or normalization was not possible due to detachment of cells, incomplete staining, or autofocus errors. For independent experiments with n=3, each biological replicate comprised at least three technical replicates. For experiments with n=2 independent observations, each observation comprised at least six technical replicates.

### Mitochondrial membrane potential measurements using TMRM

Cells were seeded the day before the assay in 96-well black-walled µClear plates (Greiner). Cells were stained with 200nM TMRM and 1µg/mL Hoechst33342 (both Life Technologies) for 30min, then medium was exchanged for inhibitor-containing fresh medium. Following 30min drug incubation, the mitochondrial membrane potential was monitored on a Cytation 5 Cell Imaging Multi-Mode Reader (Agilent Technologies) using the 4x magnification objective and the filters/LED cubes for RFP and DAPI. The TMRM mean intensity per image was normalized to the number of cells detected by the Hoechst33342 stain.

### Confocal live cell imaging

Wm3248DR cells cultured in MPM on 8-well µslides (Ibidi) were washed with HBSS^+/+^ (containing calcium and magnesium, Gibco). The cells were stained for 20 min with 1µg/ml Hoechst33342 (nuclear stain), 50nM Mitotracker green (mitochondrial stain) and 80nM TMRM (indicator of the mitochondrial membrane potential, all from Life Technologies). After washing off the staining solution with HBSS^+/+^, the cells were treated with Bam15 (2µM), FX-11 (25µM), LDH-IN-1 (15µM), or GNE140 (15µM). The signal evolution was tracked in regular intervals for 45min on the confocal Cytation 10 instrument (Agilent Technologies) using the 60x magnification objective and the filters/LED cubes for DAPI, GFP and RFP.

### Cell lysis and Western blot immunodetection

Cell lysis and Western blot analysis was performed as described before (Vollmer et al., 2009). Proteins were separated using self-cast Tris/acetate gradient gels, followed by electro-blotting onto a PVDF membrane (PVDF-PSQ, Millipore). The following antibodies were used for detection: pACC (Cell Signaling; Cat# 3661, RRID: AB_3303337), ACC (Cell Signaling; Cat# 3676, RRID: AB_2219397), p4E-BP1 (Cell Signaling; Cat# 9451, RRID:AB_330947), 4E-BP1 (Cell Signaling; Cat# 9644, RRID:AB_2097841), eIF2α (Cell Signaling; Cat# 5324, RRID: AB_10692650), GAPDH (Sigma; Cat# G9545, RRID: AB_796208). HRP-conjugated secondary antibodies were from Cell Signaling. Chemiluminescent signals were detected using a self-made ECL solution (containing 2.5mM luminol, 2.6mM hydrogen peroxide, 100mM Tris/HCl pH 8.8, and 0.2mM para-coumaric acid) (Haan & Behrmann, 2007) and a CCD camera system (Fusion FX, Vilber). Quantification was done using the Image Studio Light Software V5.2 (Li-cor). Briefly, the signals to be quantitated were normalized with respect to the non-phosphorylated protein. Each normalized signal was then divided by the mean of all normalized signals of one membrane to adjust for the possible variation of signal intensity between different membranes (each membrane including one biological replicate). The signal intensity was then represented as % of the untreated controls.

### Statistical analysis

Unless explicitly stated otherwise, three independent biological experiments were performed, each in technical triplicates. Data are presented as the means ± standard deviation (SD). Graphical representations and statistical analyses and graphical representations were performed with GraphPad Prism (version 9.5.1) software (GraphPad Software, LLC), using one-way ANOVA with Dunnett’s test correction for multiple comparisons.

## Results

### FX-11 inhibits cell proliferation of drug-sensitive and drug-resistant melanoma cells across various growth conditions

We identified FX-11 (2,3-Dihydroxy-6-methyl-7-(phenylmethyl)-4-propyl-1-naphthalenecarboxylic acid) (Figure 1A) among the best “hits” in a drug screen performed on melanoma cell lines (Preis, Rolvering, Kirchmeyer, Kreis, et al., 2024, Preprint) In this screen, FX-11 exhibited profound growth inhibitory effects in melanoma cell lines and their drug-resistant counterparts when cultivated in physiologic medium in monolayer or 3D culture systems.

**Figure 1.**
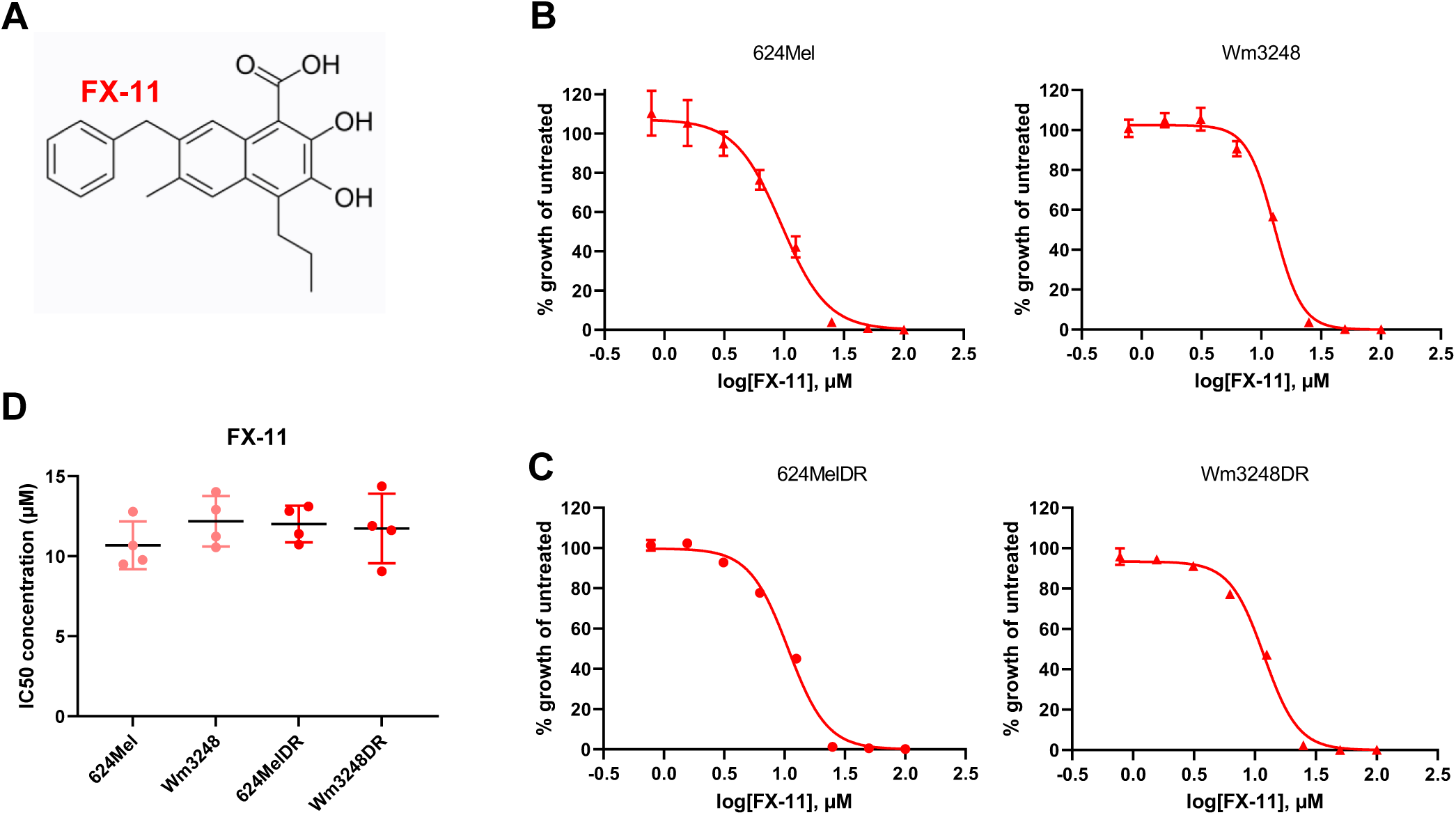
FX-11 inhibits melanoma proliferation across various growth conditions. (A) Chemical structure of FX-11 (2,3-dihydroxy-6-methyl-7-(phenylmethyl)-4-propyl-1- naphthalenecarboxylic acid). (B-C) Representative dose-response curves for FX-11 on drug-sensitive (B) and drug-resistant (C) 624Mel and Wm3248 cells grown for 72h in MPM. Mean and standard deviation of one representative biological replicate out of 4 are shown, each performed in 3 technical replicates. (D) IC_50_ values derived from the dose-response curves. Means of 4 biological replicates are shown.

Here we validated the effects of FX-11 in cell viability assays, using the BRAF^mut^ melanoma cell lines 624Mel and Wm3248 and their drug-resistant counterparts (selected by prolonged treatment with Encorafenib and Binimetinib; Preis, Rolvering, Kirchmeyer, Kreis, et al., 2024, Preprint) in monolayer cultures. Representative dose-response curves for growth in the physiologic medium MPM upon FX-11 treatment are shown in Figure 1B and 1C. The IC_50_ values are similar for drug-sensitive and resistant cell lines (Figure 1D).

### FX-11 induces cell death in melanoma cell lines

Detection of live versus dead cells upon FX-11 treatment revealed that the drug potently induced cell death of our melanoma cells (Figure 2A). To assess the possible involvement of apoptosis in cell death induction, we performed caspase 3 activity assays following drug treatment for 16h (Figure 2B) or 24h (data not shown). Staurosporine and TG101348, targeting different kinases, served as positive controls since they induce apoptosis in different cell types. Caspase 3 activity was found to be ∼5-times higher in both melanoma cell lines following FX-11 treatment as compared to untreated cells or cells treated with Encorafenib used as negative control, as it has been shown not to induce apoptosis (Z. Li et al., 2016). However, the fold induction of caspase 3 activity was lower than for the positive controls. These results indicate that apoptosis contributes at least partially to the reduced melanoma cell viability elicited by FX-11.

**Figure 2.**
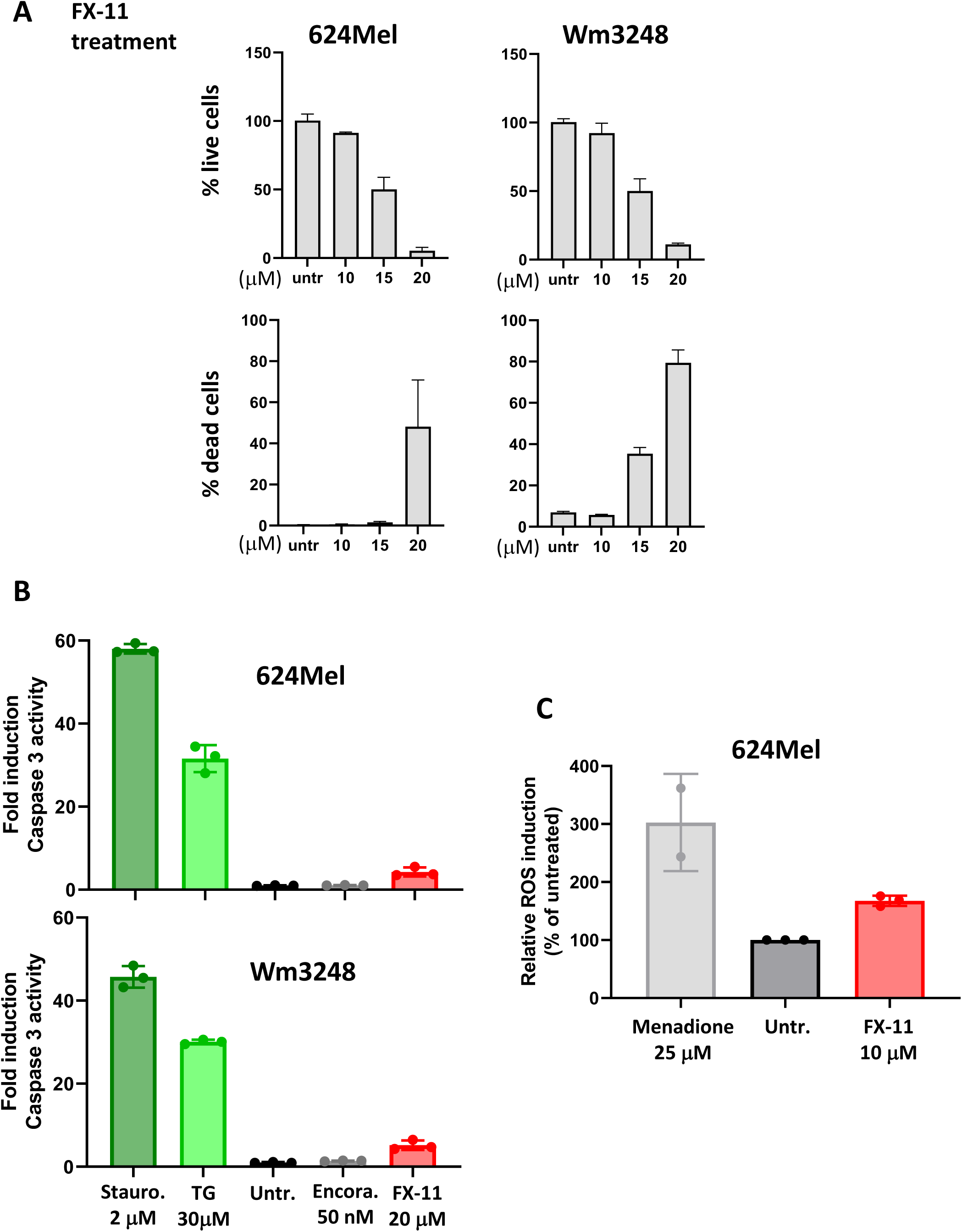
FX-11 induces cell death in melanoma cells. (A) Graphs showing the effect of different concentrations of FX-11 on cell viability in 624Mel and Wm3248 cells. After 48h of treatment, cells were stained with Hoechst 33342 and Sytox Orange for 30min and imaged on a Cytation 5. Mean and standard deviation of one representative biological replicate out of 3 are shown, each performed in 3 technical replicates, (B) Fold induction of apoptosis following 16h drug incubation as measured by Caspase 3 activity assay. Positive controls (Staurosporine (Stauro.) and TG101348 (TG)) are shown in green. Encorafenib (Encora) which leads to growth inhibition but does not induce apoptosis is also shown. Mean and standard deviation of one representative biological replicate out of 3 are shown, each performed in 3 technical replicates, (C) Relative ROS induction following 4h drug incubation of 624Mel cells. Cells were stained with CMH2DCFDA and analysed by flow cytometry in the FITC channel. Menadione serves as positive control. Means ± SD; n=3 independent experiments except for Menadione (n=2).

We further examined possible effects of FX-11 on ROS induction since exacerbated ROS can also lead to increased cell death. Cytoplasmatic ROS were quantified by flow cytometry, following a 4h drug incubation and staining with the ROS indicator CM-H_2_DCFDA. Menadione served as a positive control. We observed a ∼two-fold increased fluorescence intensity after FX-11 treatment of 624Mel cells, indicative of higher intracellular ROS levels, compared to the untreated control (Figure 2C). However, the increase in intracellular ROS levels seems to be cell-line specific: in Wm3248 cells, no increase was observed (data not shown).

### FX-11 elicits different effects compared to other LDH inhibitors in melanoma cells

The mechanism-of-action of FX-11 on cancer cells has been attributed to LDH inhibition (Le et al., 2010). As the transformation of pyruvate to lactate oxidises NADH (see Figure 3A), we measured the NAD^+^ and NADH concentrations after 24h of FX-11 exposure, in comparison to 2 other LDH inhibitors, LDH-IN-1 (Rai et al., 2017) and GNE140 (Boudreau et al., 2016), expecting to measure a decreased NAD^+^-to-NADH ratio for compounds targeting LDH. A significant decrease in the NAD^+^/NADH ratio could be observed for GNE140. For LDH-IN-1 a corresponding tendency was observed, although the result did not achieve significance in the statistical test. However, we could not detect changes in the NAD^+^/NADH ratios following FX-11 treatment (Figure 3B).

**Figure 3.**
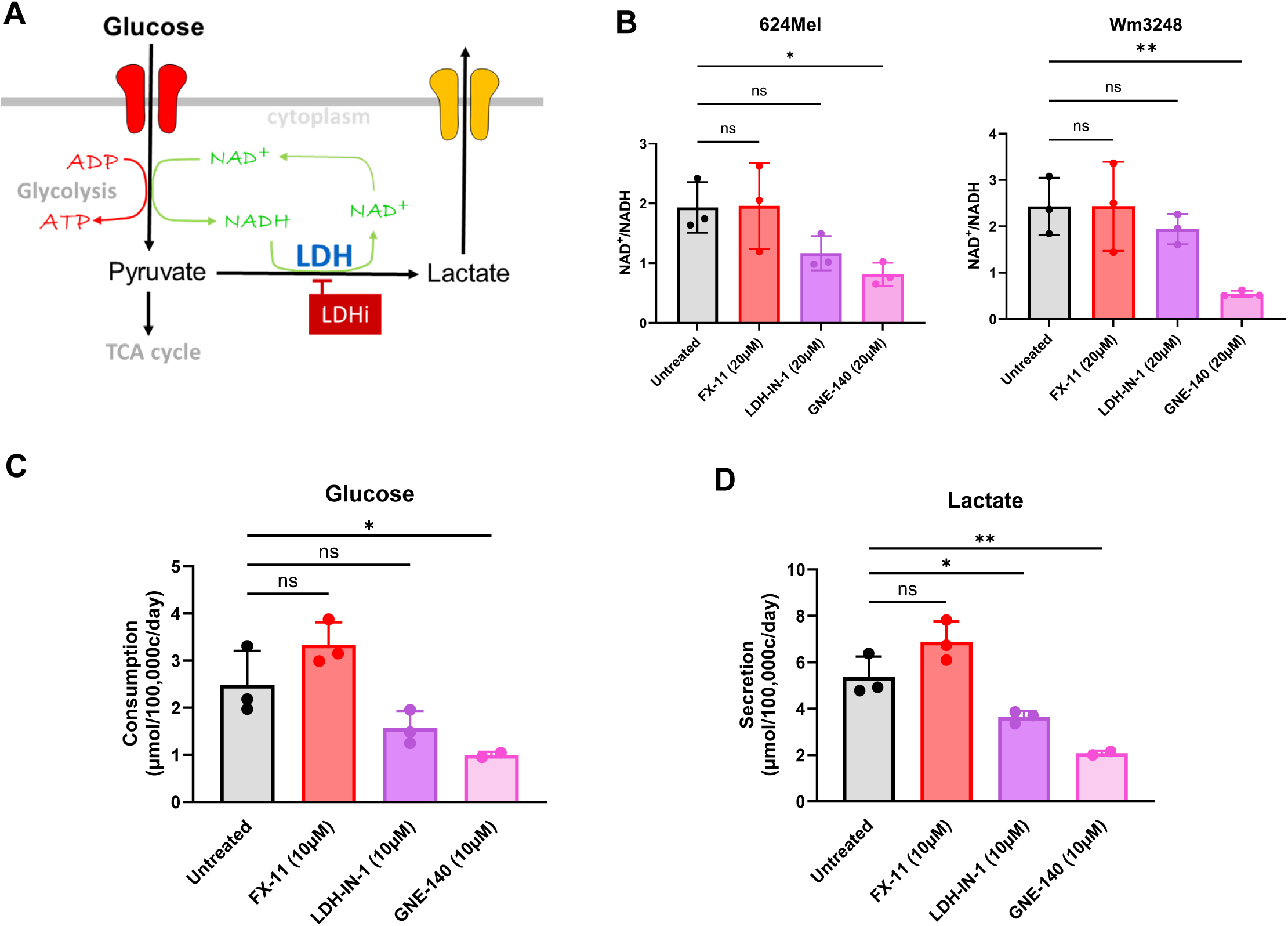
FX-11 does not recapitulate the behaviour of other LDH inhibitors in melanoma cells. (A) Scheme depicting glycolysis and the alternative fates of pyruvate: TCA cycle or lactic acid fermentation. LDH inhibition (LDHi) is highlighted in red. (B) FX-11 does not change NAD^+^/NADH ratios, as compared to the LDH inhibitors LDH-IN-1 and GNE140. (C) Glucose uptake and (D) lactate secretion in 624Mel cells grown in MPM and treated with the indicated inhibitors. Means ± SD; n=3 independent experiments, each comprising three technical replicates, except for glucose and lactate measurements for GNE140 treated cells, n=2 independent experiments. *P˂0.05, **P˂0.01.

LDH inhibition is known to suppress glycolytic activity (by suppressing NAD^+^ regeneration) and therefore to attenuate glucose uptake as well as lactate secretion of cancer cells (Boudreau et al., 2016). Consequently, we quantified glucose and lactate levels in the supernatant upon 24h of LDH inhibitor treatment (Figure 3C-D). The LDH inhibitor GNE-140 significantly reduced the glucose consumption of melanoma cells, and a corresponding tendency was seen for LDH-IN-1. Both LDH inhibitors, in addition, significantly suppressed the lactate secretion of cells. However, FX-11 did not seem to reduce the glucose uptake or the lactate secretion of cells. Instead, there was a tendency towards increased glucose consumption and lactate secretion, although the result did not achieve statistical significance. Taken together, the results indicate that FX-11 does not recapitulate the results obtained with *bona fide* LDH inhibitors in our melanoma cell lines.

### FX-11 leads to increases in OCR and ECAR, as mitochondrial uncouplers do

As the previous results were unexpected, and to investigate whether FX-11 induces metabolic alterations in melanoma cells, we evaluated the drug effects on the extracellular acidification rate (ECAR, a parameter essentially related to the glycolytic activity of cells) and the oxygen consumption rate (OCR, a measure reflecting mostly the mitochondrial respiration) using the Seahorse XFe96 Analyzer. After the initial baseline measurement of OCR and ECAR, DMSO (as solvent control) or FX-11 were injected, with concentrations of FX-11 ranging from 1 to 20µM (Figure 4A). We observe a dose- and time-dependent increase in OCR for 624Mel and Wm3248 cells at intermediate concentrations ranging from 2.5 to 10µM; at higher concentrations, OCR values decrease after the initial peak. While the increase of OCR upon FX-11 injection could be reflective of increased oxidative phosphorylation following LDH inhibition, the results from the ECAR measurements were unexpected. FX-11 injection led to a remarkable increase in the ECAR that was sustained at intermediate concentrations (Figure 4B). An LDH-inhibitor would rather be expected to decrease the ECAR, due to a decreased lactate and proton co-export. We therefore compared the response of FX-11 and other LDH inhibitors on OCR and ECAR (Figure 4C). The well characterised LDHi, LDH-IN-1 and GNE-140, led to a 20% decrease in ECAR in melanoma cells which was compensated by a 20% increase in OCR compared to baseline measurements (Figure 4D). FX-11, in contrast, led to a rise of approximately 80% in OCR and ECAR. Interestingly, a concomitant increase in OCR and ECAR, as observed for FX-11, is an effect that has been described for several mitochondrial uncouplers (Figarola et al., 2015, 2018; Kenwood et al., 2014).

**Figure 4.**
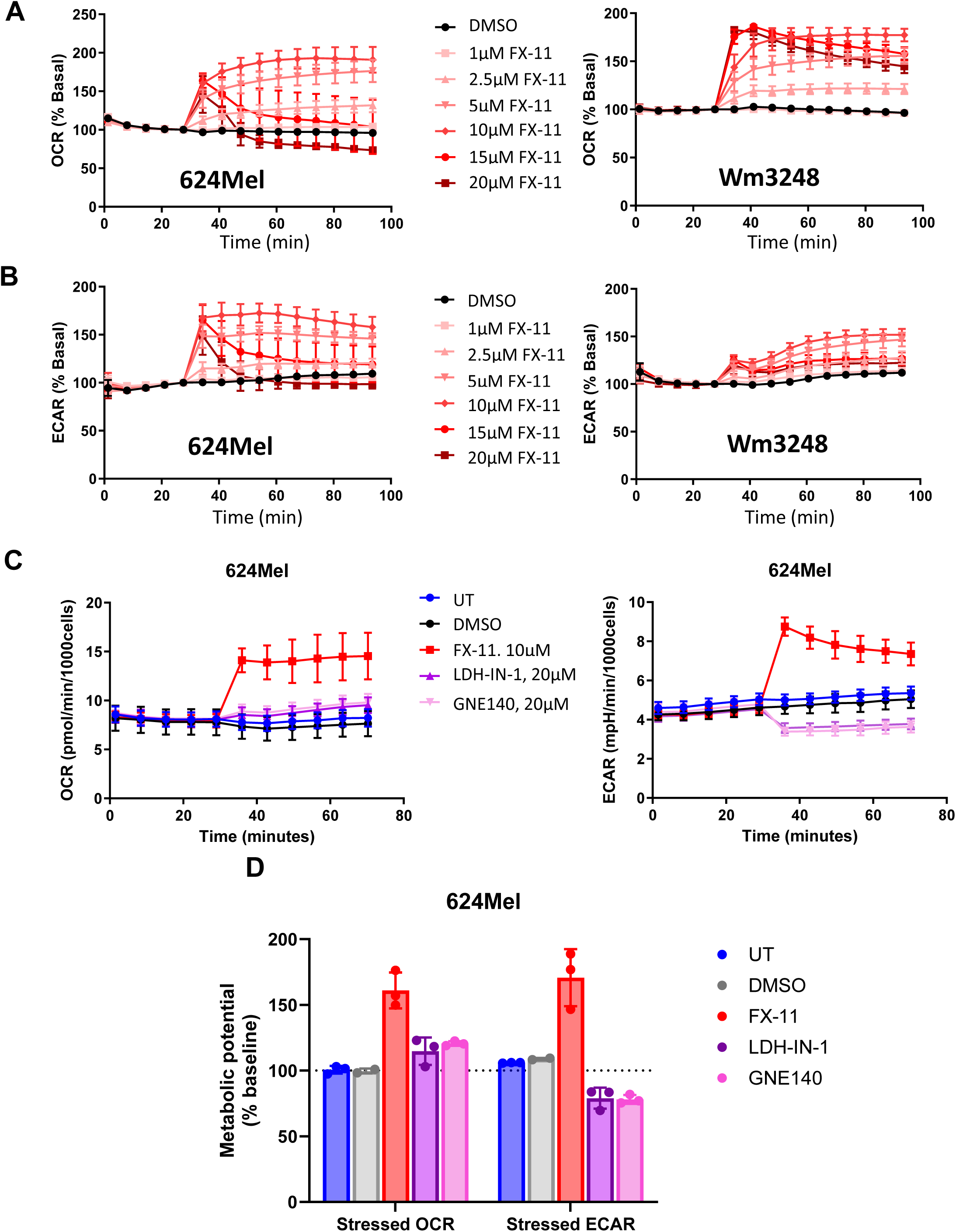
FX-11, unlike LDH inhibitors, increases OCR and ECAR of melanoma cells. (A-B) Increase in OCR (A) and in ECAR (B) following FX-11 injection at various concentrations for 624Mel and Wm3248 cells, respectively. Means ± SD from n=3 independent experiments. (C) OCR and ECAR following FX-11 or LDH inhibitor (LDH-IN-1 or GNE140) injection in 624Mel. A representative experiment of n=3 independent experiments is shown. (D) Metabolic potential (% change to basal levels) following FX-11 or LDH inhibitors injection in 624Mel (related to C). The mean ± SD of n=3 independent experiments is shown.

### FX-11 parallels the behaviour of mitochondrial uncouplers in mitochondrial stress tests

We hypothesised that FX-11 might act as a mitochondrial uncoupler. Therefore, we investigated whether FX-11 could stimulate OCR in the presence of the ATP synthase inhibitor oligomycin in the Seahorse XF Cell Mito Stress test. We compared the response of FX-11 to other well-known mitochondrial uncouplers: the prototype mitochondrial uncoupler FCCP, routinely used in the Cell Mito Stress test assay, and Bam15 (Kenwood et al., 2014). Indeed, FX-11 increased mitochondrial respiration to a similar extent as the positive controls Bam15 and FCCP, even in the presence of oligomycin (Figure 5).

**Figure 5.**
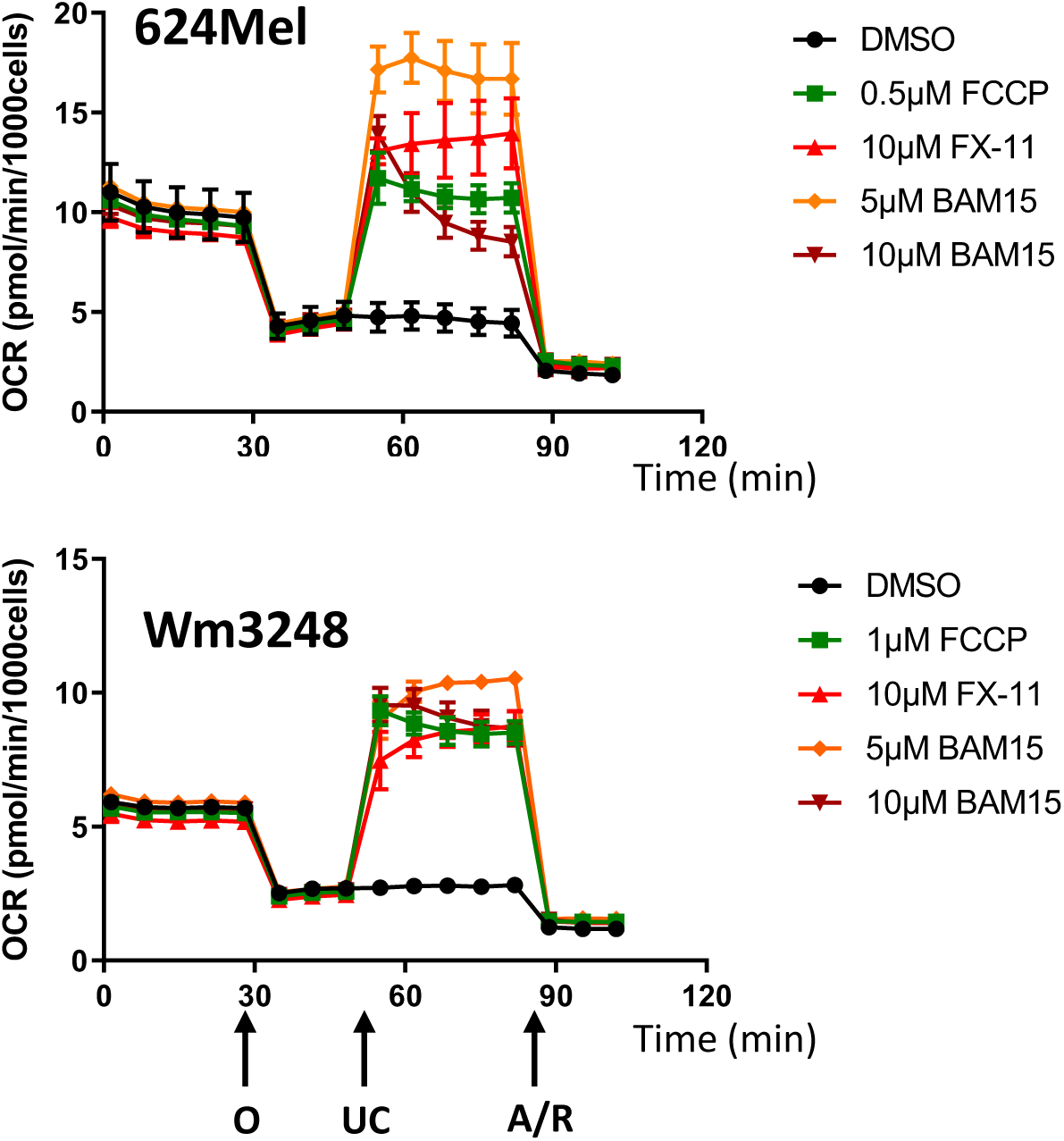
FX-11 parallels the behaviour of mitochondrial uncouplers in the mitochondrial stress test. Cell Mito stress test assay in 624Mel and Wm3248 cells. OCR after sequential injection of Oligomycin (O, 1.5µM), DMSO (ctrl) or uncoupler (UC) (FCCP, Bam15, FX-11) and Rotenone/Antimycin A (R/A; 0.5µM). Representative experiment of n=2 independent experiments. For the representation, values from at least six technical replicates per condition are shown.

### FX-11 affects cellular energy homeostasis through AMPK-mTOR signalling

Since mitochondrial uncouplers (as well as LDH inhibitors) should lead to a reduction of ATP levels and of the ATP/ADP ratio, which ultimately results in activation of AMPK (Shrestha et al., 2021), we performed Western blot analyses to investigate phosphorylation events downstream of AMPK. Also here, FCCP and Bam15 were used as positive controls. AMPK is known to inhibit Acetyl-CoA Carboxylase (ACC) by phosphorylation and is also described to inhibit mammalian target of rapamycin (mTOR) (Figure 6A). 624Mel responded to FCCP, Bam15, and FX-11 with increased phosphorylation of ACC (Figure 6B and C). Impairment of mTOR activity was indicated by a decrease in 4E-BP phosphorylation, a direct downstream target of mTOR (Figure 6B and C).

**Figure 6.**
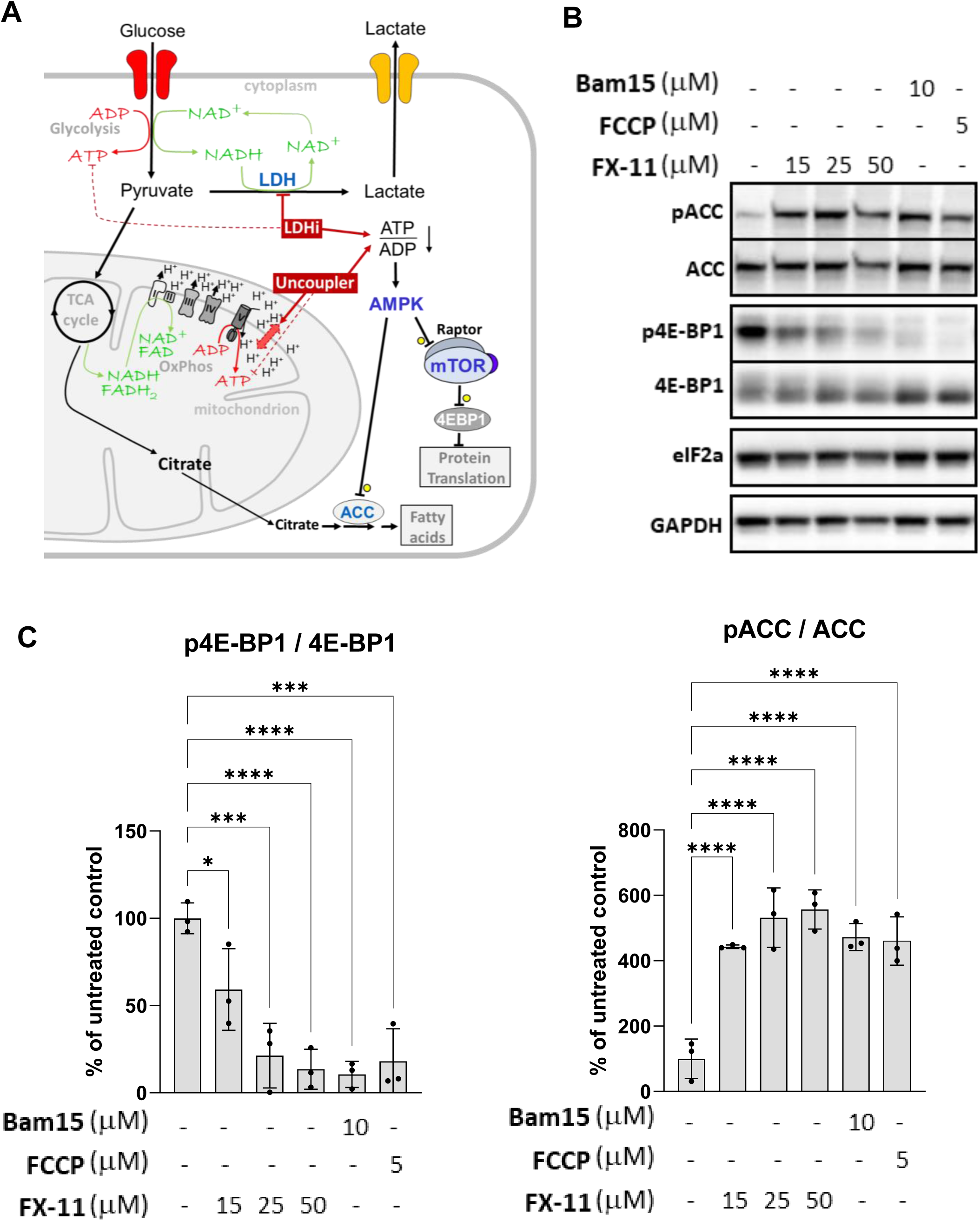
FX-11 affects cellular energy homeostasis through AMPK-mTOR signalling. (A) Scheme depicting the impact of LDH inhibitors and of uncouplers on the cellular energy metabolism. (B) 624Mel cells were treated for 6h with the indicated concentrations of FX-11 or with FCCP (5µM) or Bam15 (10µM). Lysates of cells were separated by SDS PAGE. Western blots were detected with antibodies against phospho-ACC (pACC) and phospho-4E-BP1 (p4E- BP1). After stripping of the membranes, the blots were re-detected with antibodies against total ACC, eIF2a, 4E-BP1, and GAPDH as loading controls. (C) Mean and standard deviation of three biological replicates of Western blot results shown in (B). The signals were normalised with respect to the untreated control signal which is represented as 100%. *P˂0.05, **P˂0.01, ***P˂0.001, ****P˂0.0001.

However, as compared to FCCP and Bam15, higher concentrations of FX-11 were required to achieve the same effects.

Taken together, our results indicate that FX-11 affects the energy metabolism of melanoma cells and activates AMPK, which in turn inhibits mTOR activity.

### FX-11 impairs the mitochondrial membrane potential

Mitochondrial uncouplers dissipate the proton gradient across the inner mitochondrial membrane (Figure 6A). Thus, we measured the mitochondrial membrane potential (ΔΨ_M_) by TMRM staining and quantified the fluorescence signal after 30min of drug incubation. Increased proton leak into the mitochondrial matrix due to uncoupling is expected to induce mitochondrial depolarization, reflected by a reduction in TMRM intensity. We included the mitochondrial uncouplers FCCP and Bam15 as positive, and LDH-IN-1 as negative control. In both 624Mel (Figure 7A) and Wm3248 cells (Figure 7B), FX-11 caused a rapid decrease in the mitochondrial membrane potential, comparable to the decrease observed for FCCP and Bam15. As expected, we did not observe a drastic impairment of the TMRM staining upon treatment with LDH-IN-1, although it caused a small decrease in the mitochondrial membrane potential in Wm3248 cells.

**Figure 7:**
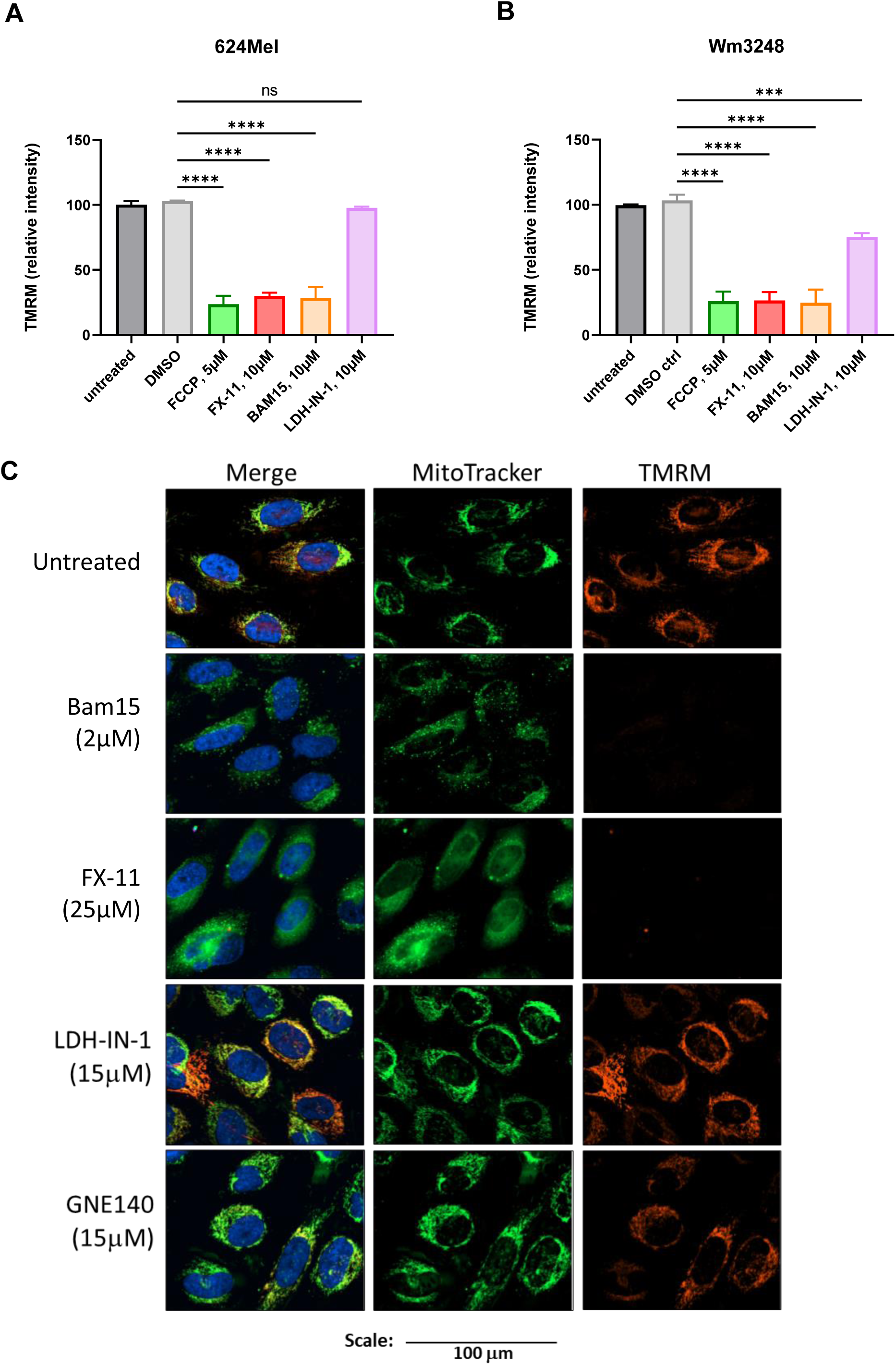
FX-11 profoundly affects the mitochondrial membrane potential. (A) 624Mel and (B) Wm3248 cells were treated with the indicated concentrations of FCCP, FX-11, Bam15, or LDH-IN-1 for 30min. The measured TMRM intensity was normalized to the DMSO control, n=3 independent experiments, mean and standard deviation are shown. (C) Wm3248DR cells were stained with Hoechst33342 (blue, nucleus), TMRM (red, indicates the mitochondrial membrane potential), and Mitotracker Green (green, mitochondria), and treated with Bam15, FX-11, or LDH inhibitors (LDH-IN-1, GNE140) at the indicated concentrations for 20 min. Confocal images were taken with a Cytation 10 instrument. Scale: 100µm. ***P<0.001, ****P<0.0001.

We also investigated mitochondrial appearance upon drug treatment by confocal microscopy. Cells were stained with Mitotracker Green, TMRM, and Hoechst33342 (nuclear stain), treated with drugs, and imaged at 60X magnification. Representative confocal images of mitochondrial staining with TMRM and Mitotracker Green in Wm3248DR cells following 20min drug incubation are shown (Figure 7C). The LDH inhibitors, LDH-IN-1 and GNE140, do not show differences in the aspect of the mitochondrial stains. Bam15, however, very efficiently reduced TMRM intensity, which was no longer observable after 5min of administration of the drug (data not shown). Also, the localisation of Mitotracker Green was drastically changed upon Bam15 treatment. It was diffuse at early time points (not shown), followed by a punctuate distribution. Like Bam15 treatment, FX-11 reduced TMRM intensity and induced adiffuse localisation of Mitotracker Green, and punctuate structures appeared. Of note, the concentration of FX-11 was more elevated than the one of Bam15, the effects induced were slower to develop and less drastic, emphasising the high efficiency of Bam15 as mitochondrial uncoupler.

## Discussion

Tumour cells often have a higher dependency on glycolysis, so that inhibitors of glycolytic enzymes including lactate dehydrogenase are regarded as promising drug candidates (Claps et al., 2022; Sharma et al., 2022). LDH is overexpressed in many cancers, and high levels of the enzyme correlate with poor patient survival (Cheng et al., 2021; Claps et al., 2022). For instance, *BRAF*-mutated melanoma cells were reported to have upregulated glycolysis (Abildgaard & Guldberg, 2015). Treatment with targeted therapy against mutated BRAF and the downstream kinase MEK leads to an initial downregulation of glycolysis as a consequence of MAPK pathway inhibition. However, drug-resistant melanoma cells restore glycolysis after an initial response to the MAPKi and show an increased oxidative metabolism (Alkaraki et al., 2021). Our drug-resistant melanoma cell lines used in this study recapitulate this behaviour (Preis, Rolvering, Kirchmeyer, Kreis, et al., 2024, Preprint).

FX-11 is effective in various cancer models, *in vitro* and *in vivo*, alone or in combination with other drugs (Gao et al., 2016; Hou et al., 2021; Kim et al., 2019; Mohammad et al., 2019; Rajeshkumar et al., 2015; Xian et al., 2015; Xu et al., 2019). FX-11 was described as a NADH-competitive LDH inhibitor and had growth inhibitory effects on lymphoma, pancreatic cancer, renal cell carcinoma, and breast cancer cells (Le et al., 2010). In several studies, FX-11 was found to be particularly effective in tumours relying on glycolysis (Le et al., 2010; Rajeshkumar et al., 2015; Xu et al., 2019).

We identified FX-11 as one of the promising chemical compounds from a pharmacological screen with a metabolism-oriented library using physiologic medium in monolayer and 3D cell culture models of drug-sensitive and drug-resistant melanoma cells (Preis, Rolvering, Kirchmeyer, Kreis, et al., 2024, Preprint). The validation experiments presented here confirmed that FX-11 was effective, both in drug-sensitive and -resistant melanoma cell lines (Figure 1). Considering the above-mentioned differences in metabolism reported for *BRAF*-mutated drug-naive and -resistant cells, it is noteworthy to mention that the IC_50_ values were similar in the drug-sensitive and - resistant melanoma cell lines (Figure 1D).

When performing experiments to validate the mechanism-of-action of FX-11 as LDH inhibitor, we observed effects that were difficult to reconcile with the described function. While the LDH inhibitors used as positive controls (LDH-IN-1, GNE140) led to a reduction in the NAD^+^/NADH ratio in cells, with a reduction of glucose consumption and lactate secretion, FX-11 treatment failed to recapitulate the expected effects (Figure 3).

As our results seem contradictory to previous findings (Le et al., 2010), we made use of the Seahorse metabolic analyser to capture metabolic alterations upon FX-11 injection in real time. Cells treated with LDH inhibitors show a reduced lactated production and secretion (in co - transport with protons), thus one would expect a reduction of ECAR (Oshima et al., 2020). However, FX-11 treatment induced a strong increase of the ECAR as well as of the OCR (Figure 4). This result was unexpected and reminiscent of mitochondrial uncouplers: Mitochondrial uncouplers induce a proton leak across the inner mitochondrial membrane into the matrix (Rawling et al., 2020), leading to futile cycles of nutrient oxidation in the TCA cycle without concomitant ATP production, which is the basis for their long-known effect as weight loss drugs (Childress et al., 2018). The elevated amounts of the reduced redox cofactors NADH and FADH_2_ intensify the electron flow through the electron transport chain to O _2_, thereby leading to an increased OCR. To further support the hypothesis of FX-11 working as an uncoupler, we performed the mitochondrial stress test on the Seahorse and included the *bona fide* uncouplers (Bam15, and FCCP) as positive controls. FX-11, as well as the well-known uncouplers, were able to induce the OCR of cells, even in the presence of the ATP synthase inhibitor Oligomycin (Figure 5).

We also found that FX-11 modulated the AMPK/mTOR signalling pathway in melanoma cells (Figure 6), in accordance with previous results (Hou et al., 2021; Le et al., 2010). While this effect has been attributed to LDH inhibition by FX-11 (Cheng et al., 2021), it has also been previously described for mitochondrial uncouplers (Kenwood et al., 2014). Indeed, we provide evidence that uncouplers activate AMPK and inhibit mTOR, by investigating their downstream signalling events (phosphorylation of ACC for AMPK and dephosphorylation of 4E-BP1 for mTOR).

Lastly, we find FX-11 to rapidly dissipate the mitochondrial membrane potential, a mechanism characteristic for mitochondrial uncouplers, such as Bam15, while we saw no or only minor effects of LDH inhibitors on TMRM staining (Figure 7). Interestingly, also the intensity of MitoTracker Green was reduced with Bam15, which is probably explained by the fact that this dye may also depend on the mitochondrial potential but is less sensitive to depolarisation than TMRM.

Taken together, our comparison of FX-11 to *bona fide* uncouplers (Bam15, FCCP) and LDH inhibitors (LDH-IN-1, GNE140) revealed that the effects observed for FX-11 rather recapitulate those of uncouplers in our melanoma lines: (i) an LDH inhibitor would be expected to attenuate the lactate accumulation in the medium and to inhibit the glucose uptake, which was not the case for FX-11. Moreover, FX-11 did not affect the NAD^+^/NADH ratio. (ii) Like known uncouplers (and unlike known LDHi), FX-11 increased both the OCR and the ECAR. (iii) FX-11 caused a rapid decrease in mitochondrial membrane potential (ΔΨ_m_), an effect we did not observe for LDH inhibitors. Therefore, we propose a different mechanism-of-action for FX-11.

We are not the first group to describe effects for FX-11 which do not support the function of FX- 11 as an LDH inhibitor. Similar to our results, in MDA-MB-230 breast cancer cells, FX-11 also induced an increased OCR, an increased ECAR, and increased secretion of lactate which is not expected for a LDH inhibitor (Jiang et al., 2021). Interestingly, we also observed a tendency of increased lactate secretion and increased glucose consumption upon FX-11 treatment (Figure 3C), although these results did not achieve statistical relevance in our tests. These and the results of others (Jiang et al., 2021) could, however, be explained by increased reliance on glycolysis for ATP production when the mitochondrial ATP production is hampered due to dissipation of the proton gradient. To our knowledge we are the first to provide evidence for FX-11 to work as a mitochondrial uncoupler in cancer cells. Moreover, FX-11 contains an acid group attached to a substituted delocalised naphthalene (Figure 1A), a molecular feature typical for uncouplers which often present acidic groups within delocalised π-systems (Childress et al., 2018; Rawling et al., 2020).

Interestingly, other mitochondrial uncouplers have shown activity in some experimental models of melanoma (Figarola et al., 2018; Y. Li et al., 2014; Serasinghe et al., 2018). The chemical uncouplers SR4 and niclosamide reduced proliferation of various melanoma cells (Figarola et al., 2018). Both uncouplers activated AMPK signalling and inhibited mTOR signalling, similar to the findings in the present study. Importantly, both uncouplers were also effective in mouse xenografts with no overt side effects in other organs. It was found that melanoma cells with a stronger OXPHOS phenotype, with a mutation in the *LKB1* gene, or with acquired resistance to the BRAF inhibitor vemurafenib displayed a greater sensitivity to both uncouplers (Figarola et al., 2018). However, we did not observe a strong increase in the sensitivity when comparing parental cells and their derivates resistant against the combination of Encorafenib and Binimetinib. Serasinghe et al. investigated Bam15 in the context of targeted therapy (Vemurafenib/Trametinib) against melanoma cells. While they did not observe any toxicity of Bam15 alone (i.e., no effects on proliferation, cell death or colony formation), the combination with either Vemurafenib/Trametinib and/or a BH3-mimetic strongly promoted apoptosis (Serasinghe et al., 2018). Bam15 has been described to have a broad range of efficacy with low toxicity (Kenwood et al., 2014; Serasinghe et al., 2018). Bam15 specifically uncouples mitochondrial membranes and, in contrast to many other uncouplers, has low activity on other cellular membranes (Kenwood et al., 2014).

While we could show that FX-11 acts as a mitochondrial uncoupler rather than an LDHi in our melanoma cells, the spectrum of effects of FX-11 warrants further investigation. However, because it is used at relative high concentrations (10-20µM), drug toxicity and off-target effects may limit its applicability. Bam15, having a similar mechanism as FX-11, shows higher potency, selectivity for uncoupling of mitochondrial membranes, and has low toxicity (Kenwood et al., 2014; Serasinghe et al., 2018). In melanoma cells resistant to targeted therapy, it might be especially interesting to combine such low toxicity compounds with other treatments which, e. g., inhibit autophagy, induce further mitochondrial stress, or inhibit glycolysis.

## Acknowledgements

The 624Mel were a kind gift of Dr. Ruth Halaban (Dermatology Department, Yale School of Medicine, USA). We thank the Luxembourg Centre for Systems Biomedicine (LCSB) Metabolomics Platform for running metabolomics samples (YSI measurements to determine glucose and lactate concentrations).This work was supported by the Luxembourg National Research Fund (FNR), project number 10675146 (funding scheme PRIDE, “CANBIO”), and by the Fondation Cancer Luxembourg (“SecMelPro” grant).

## Abbreviations

ACC: Acetyl-CoA carboxylase
AMPK: AMP-activated kinase
DMSO: dimethyl sulfoxide
DR: drug-resistant
ECAR: extracellular acidification rate
eIF2α: eukaryotic initiation factor 2α
FITC: fluorescein isothiocyanate
GAPDH: glyceraldehyde 3-phosphate dehydrogenase
GFP: green fluorescent protein
LDH: lactate dehydrogenase
MAPK: mitogen activated protein kinase
MPM: modular physiologic medium
mTOR: mammalian target of rapamycin
OCR: oxygen consumption rate
TMRM: tetramethyl rhodamine methyl ester
4E-BP1: 4E-binding protein 1
ROS: reactive oxygen species

